# The Mus musculus papillomavirus type 1 E7 protein binds to the retinoblastoma tumor suppressor - implications for viral pathogenesis

**DOI:** 10.1101/2021.07.30.454561

**Authors:** Tao Wei, Miranda Grace, Aayushi Uberoi, James C. Romero-Masters, Denis Lee, Paul F. Lambert, Karl Munger

## Abstract

The species specificity of papillomaviruses has been a significant roadblock for performing in vivo pathogenesis studies in common model organisms. The Mus musculus papillomavirus type 1 (MmuPV1) causes cutaneous papillomas that can progress to squamous cell carcinomas in laboratory mice. The papillomavirus E6 and E7 genes encode proteins that establish and maintain a cellular milieu that allows for viral genome synthesis and viral progeny synthesis in growth-arrested, terminally differentiated keratinocytes. The E6 and E7 proteins provide this activity by binding to and functionally reprogramming key cellular regulatory proteins. The MmuPV1 E7 protein lacks the canonical LXCXE motif that mediates the binding of multiple viral oncoproteins to the cellular retinoblastoma tumor suppressor protein, RB1. Our proteomic experiments, however, revealed that MmuPV1 E7 still interacts specifically with RB1. We show that MmuPV1 E7 interacts through its C-terminus with the C-terminal domain of RB1. Binding of MmuPV1 E7 to RB1 did not cause significant activation of E2F-regulated cellular genes. MmuPV1 E7 expression was shown to be essential for papilloma formation. Experimental infection of mice with MmuPV1 virus expressing an E7 mutant that is defective for binding to RB1 caused delayed onset, lower incidence, and smaller sizes of papillomas. Our results demonstrate that the MmuPV1 E7 gene is essential and that targeting non-canonical activities of RB1, which are independent of RB1’s ability to modulate the expression of E2F-regulated genes, contribute to papillomavirus-mediated pathogenesis.

**Importance:** Papillomavirus infections cause a variety of epithelial hyperplastic lesions, warts. While most warts are benign, some papillomaviruses cause lesions that can progress to squamous cell carcinomas and approximately 5% of all human cancers are caused by human papillomavirus (HPV) infections. The papillomavirus E6 and E7 proteins are thought to function to reprogram host epithelial cells to enable viral genome replication in terminally differentiated, normally growth-arrested cells. E6 and E7 lack enzymatic activities and function by interacting and functionally altering host cell regulatory proteins. Many cellular proteins that can interact with E6 and E7 have been identified, but the biological relevance of these interactions for viral pathogenesis has not been determined. This is because papillomaviruses are species-specific and do not infect heterologous hosts. Here we use a recently established mouse papillomavirus (MmuPV1) model to investigate the role of the E7 protein in viral pathogenesis. We show that MmuPV1 E7 is necessary for papilloma formation. The retinoblastoma tumor suppressor protein (RB1) is targeted by many papillomaviral E7 proteins, including cancer-associated HPVs. We show that MmuPV1 E7 can bind RB1 and that infection with a mutant MmuPV1 virus that expresses an RB1 binding defective E7 mutant caused smaller and fewer papillomas that arise with delayed kinetics.

## INTRODUCTION

Papillomaviruses (PVs) have been isolated from a wide range of vertebrate species. They have a tropism for squamous epithelia, and individual genotypes often have a marked preference for infecting mucosal or cutaneous squamous epithelia. Approximately 440 human PVs (HPVs) have been identified and they are phylogenetically classified into several genera (1). Among these, most alpha genus HPVs preferentially infect mucosal epithelia. A group of approximately 15 “high-risk” alpha genus HPVs are the etiological agents of almost all cervical cancers and a large percentage of other anogenital tract carcinomas as well as a growing fraction of head and neck squamous cell carcinomas (SCCs), particularly oropharyngeal cancers (2, 3). Overall, high-risk HPV infections contribute to >5% of all human cancers (4). The beta and gamma genus HPVs mostly infect cutaneous epithelia and infections with some of these HPVs contribute to the development of cutaneous squamous cell carcinomas (cSCCs) in individuals afflicted by the rare hereditary disease, epidermodysplasia verruciformis (5, 6), or in long-term systemically immune-suppressed organ transplant patients (7–9).

Cell- and animal model-based studies have revealed that the E6 and E7 proteins of high-risk alpha, as well as the cSCC-associated beta HPVs, have oncogenic activities. HPV E6 and E7 encode small, cysteine-rich, metal-binding proteins. They lack intrinsic enzymatic activities, do not function as DNA binding transcription factors, and do not share extensive sequence similarities with cellular proteins. By binding to and interfering with the functionality of important, host regulatory proteins they elicit profound alterations in cellular physiology to permit long-term viral persistence as well as viral progeny synthesis (10, 11). In rare cases, these alterations and the cellular responses that are triggered can cause cancer formation (12). A large number of potential cellular protein targets have been identified for the high-risk alpha HPV E6 and E7 proteins through proteomic studies (10, 11). Similar proteomic experiments with the beta HPV E6 and E7 proteins have revealed that, while they share some interactors with high-risk mucosal HPVs, they also interact with distinct cellular proteins and signaling pathways (13).

Papillomaviruses are species-specific and cannot productively infect heterologous host organisms. Hence the biological relevance of specific interactions of the HPV E6 and E7 proteins with specific host pathways cannot be tested in infectious animal models. Traditional animal models of PV infection and pathogenesis are limited to species that are not genetically tractable. The discovery of the Mus musculus papillomavirus type 1 (MmuPV1), which can be used to experimentally infect standard laboratory mice, has finally provided a viable experimental model system to explore the importance of specific virus-host interactions in viral pathogenesis. MmuPV1 was discovered based on its ability to cause cutaneous papillomas and provides us an opportunity to better understand molecular mechanisms by which HPVs promote cutaneous disease in an *in vivo* animal model (14).

We have previously reported that the MmuPV1 E6 protein shares with the beta HPV8 E6 protein the ability to inhibit NOTCH and TGF-beta signaling by interacting with the NOTCH co-activator MAML and the DNA binding SMAD2 and SMAD3 proteins that are downstream of TGF-beta signaling (15). Moreover, by experimentally infecting with MmuPV1 mutant genomes we showed that the presence of a functional MAML1 binding site on E6 is critical for papilloma formation (15). We have now extended these proteomic studies to the MmuPV1 E7 protein. Here we show that MmuPV1 E7 expression is necessary for papilloma formation. Like some gamma HPV E7 proteins, MmuPV1 E7 lacks an LXCXE (L, leucine; C, cysteine; E, glutamic acid; X, any amino acid)-based binding site for the retinoblastoma tumor suppressor, RB1 (16, 17). Using affinity purification of MmuPV1 E7 associated cellular protein complexes followed by mass spectrometry, we discovered that MmuPV1 E7 can bind RB1. Similar to some animal PV and gamma HPV E7 proteins that also bind RB1 despite lacking LXCXE domains, the RB1 binding site maps to the MmuPV1 E7 C-terminus. Experimental MmuPV1 infection of a mouse strain that expresses an LXCXE protein binding deficient RB1 mutant, causes the formation of papillomas. This result shows that the ability of MmuPV1 to cause papillomas is not dependent on binding RB1 through its LXCXE binding cleft. Consistent with this finding, we mapped the MmuPV1 E7 binding site to the RB1 C-terminus. Unlike LXCXE containing E7 proteins, MmuPV1 E7 expression did not trigger efficient activation of E2F-responsive cellular genes. Experimental infection with a MmuPV1 mutant virus that encodes an RB1 binding defective E7 protein, inefficiently caused papillomas when compared to the wild type virus, and the lesions that did arise were smaller and appeared later than those arising in wild type MmuPV1 infected animals. These findings support the hypothesis that MmuPV1 E7 contributes to pathogenesis, at least in part, through its interactions with RB1.

## RESULTS

### MmuPV1 E7 is required for viral pathogenesis

To understand how MmuPV1 mediates its pathogenesis, we first asked whether the viral E7 gene is required for the virus to induce papillomas. To address this question, we engineered a stop codon immediately after the ATG initiation codon in the MmuPV1 E7 translational open reading frame in the context of the full-length MmuPV1 DNA genome. MmuPV1 E7 only contains one translational start codon, so no E7-related polypeptides can be expressed from internal methionine residues. Nor have there been identified any spliced MmuPV1 mRNAs that could produce E7-related polypeptide initiating from a start codon present in an upstream open reading frame (18). MmuPV1 quasivirus containing the wild type or E7 stop mutant were generated in vitro in 293FT cells as described previously (19). The yield of virus particles containing encapsidated viral genomes was determined by quantifying the amount of DNAse resistant viral genomes in the fractions from the density gradient (19). These virus particles are referred to as “quasiviruses” to distinguish them from authentic viruses generated in naturally infected tissue. The infectivity of these viral stocks was confirmed by RT-PCR detection of viral E1^E4 spliced mRNAs expressed at 48 hours after infecting cultured mouse keratinocytes **(Figure 1A)**.

**Figure 1.**
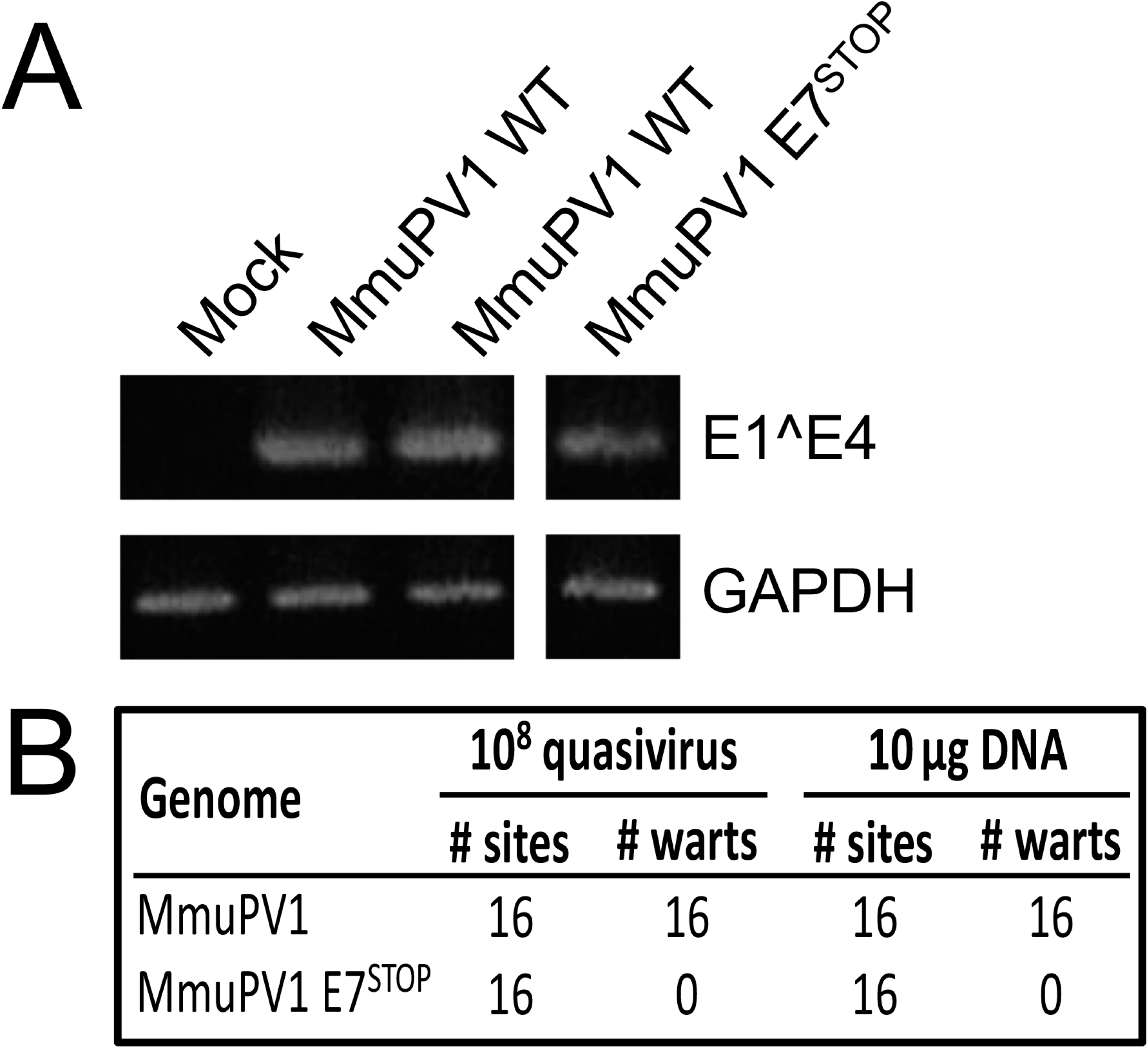
MmuPV1 lacking E7 expression does not induce warts. The MmuPV1 E7^STOP^ quasivirus is infectious. To assess infectivity of quasivirus stocks, mouse JB6 keratinocytes were exposed to quasivirus and 48 hours later RNA was extracted and subjected to reverse transcription-coupled polymerase chain reaction to detect MmuPV1 E1^E4 transcripts (top panel). GAPDH expression is shown as a control (bottom panel). Samples were run on the same gel, with irrelevant lanes in the middle cropped out in **(A)**. Tails and ears of nude mice were scarified and infected with the indicated amounts of quasiviruses or DNA and monitored for wart formation over 3 months. Neither MmuPV1 E7^STOP^ quasivirus nor MmuPV1 E7^STOP^ DNA induced wart formation, while wild type MmuPV1 quasivirus or DNA induced warts with a 100% penetrance **(B)**.

Stocks of quasiviruses containing either wild type (MmuPV1) or E7-null (MmuPV1 E7^STOP^) genomes were used to infect Nude-FoxN1^*nu/nu*^ mice at cutaneous sites, both on the ears and tail, at the same dose of 10^8^ viral genome equivalents (VGE) that was shown previously to induce papillomas at 100% of sites infected with wild type MmuPV1 (20). Briefly, sites were wounded by lightly scarifying the epidermis with a needle, then a solution containing quasivirus was applied to the wounded skin. Mice were monitored for papilloma formation weekly for 3 months. As observed before (20), wild-type MmuPV1 quasivirus caused papilloma formation at 100% of infected sites on the nude mice, whereas infections with the mutant MmuPV1 E7^STOP^ quasivirus did not induce any papillomas (**Figure 1B**). Mock-infected nude mice also did not develop papillomas (15). The experiment was repeated by placing on wounded sites naked re-circularized viral genomes, which have been reported to be infectious and cause papilloma formation (13, 21). Consistent with our previous findings (15), 10 μg of re-circularized wild type MmuPV1 genome caused papillomas at 100% of sites exposed to the viral DNA on the nude mice by the end of 3 months post-infection, whereas 10 μg of re-circularized MmuPV1 E7^STOP^ viral DNA failed to induce papillomas at any sites **(Figure 1B)**. Together, these results indicate that the expression of the viral E7 protein is required for MmuPV1 to induce papillomatosis *in vivo*.

### MmuPV1 E7 lacks an LXCXE motif but can bind to RB1

The retinoblastoma tumor suppressor, RB1, is an important cellular target of many PV E7 proteins. Most PV E7 proteins interact with RB1 through a conserved, N-terminal LXCXE motif (11). The MmuPV1 E7 protein, however, lacks an LXCXE sequence (**Figure 2A**). Some PVs including the canine papillomavirus type 2 (CPV2) and the gamma HPV4 and HPV197 E7 proteins have been shown to bind RB1 despite lacking LXCXE domains (16, 17). To determine whether MmuPV1 E7 may bind RB1 we performed affinity purification/mass spectrometry (AP/MS) experiments. N-terminal and C-terminal FLAG/HA epitope-tagged MmuPV1 E7 proteins were transiently expressed in HCT116 human colon carcinoma cells. HCT116 cells are used because they are of epithelial origin, are highly transfectable, express wild-type RB1 and do not contain any known exogenous viral sequences. MmuPV1 E7-associated proteins complexes were isolated by affinity chromatography on HA antibody resin, eluted with HA peptide, and analyzed by mass spectrometry. These experiments revealed that MmuPV1 E7 can interact with RB1 as evidenced by the detection of 34 and 16 unique RB1 peptides in experiments performed with N-terminally and C-terminally tagged MmuPV1 E7, respectively (Table S1). We confirmed the MmuPV1 E7/RB1 interaction by transfecting HCT116 cells with an expression vector encoding N-terminally FLAG/HA-tagged MmuPV1 E7 followed by immunoprecipitation/western blot analysis (**Figure 2B**). We also found MmuPV1 E7 to bind to endogenous murine Rb1 in similar IP/western experiments performed in mouse NIH3T3 fibroblasts using transfected N-terminally FLAG/HA-tagged MmuPV1 E7 (**Figure 2C**). That MmuPV1 E7 can bind both human and murine retinoblastoma proteins is not surprising; they are highly conserved. Because of a lack of availability of expression vectors for murine Rb1, subsequent studies characterizing the nature of this interaction, described below, were necessarily carried out using expression vectors for wild-type or mutant forms of human RB1.

**Figure 2.**
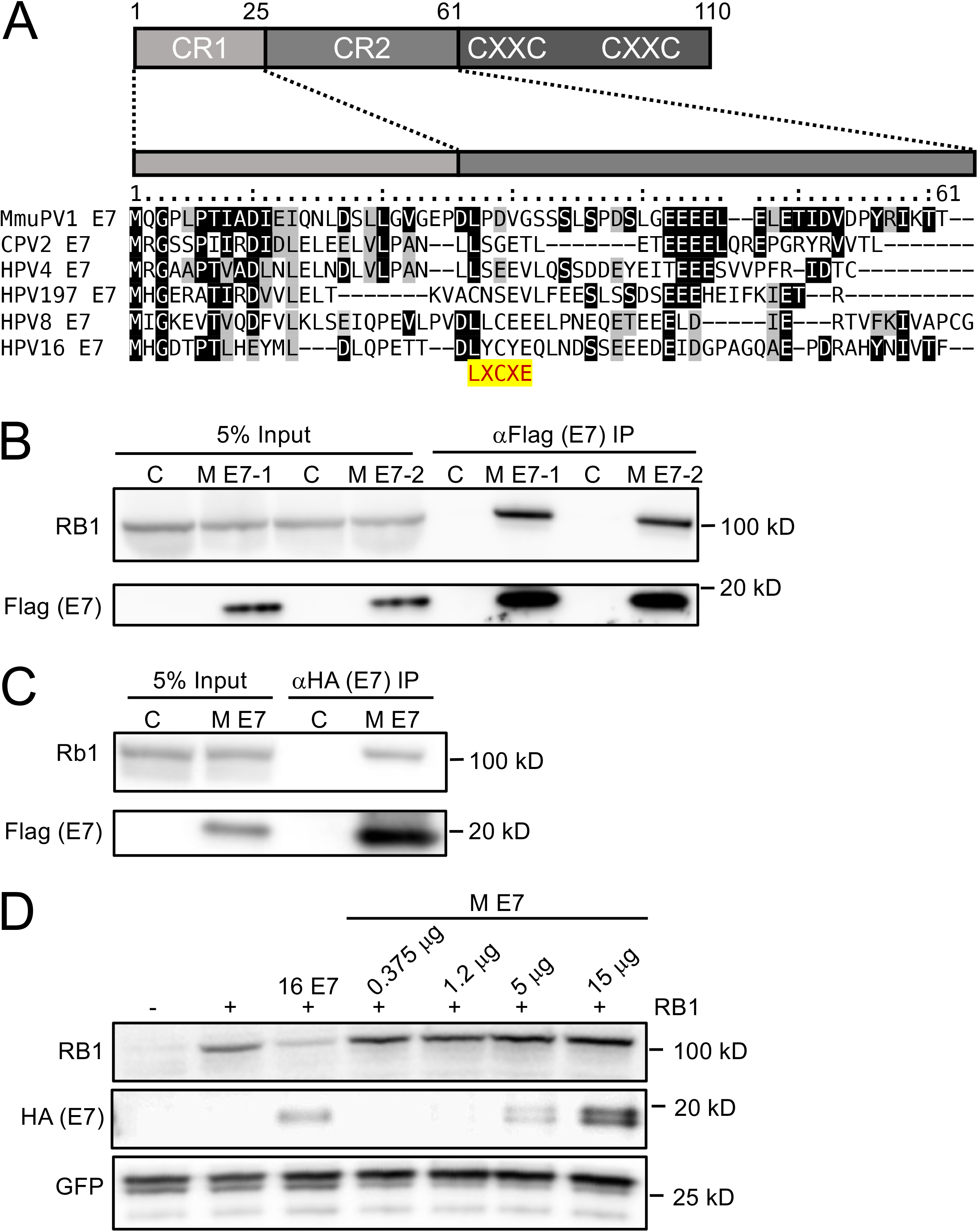
MmuPV1 E7 binds but does not destabilize the retinoblastoma tumor suppressor protein, RB1. Alignment of the N-terminal sequences of MmuPV1 E7 and the canine papillomavirus 2 (CPV2), γ1-HPV4, γ24-HPV197, β1-HPV8, and α9-HPV16. Identical residues are marked by black boxes, chemically similar residues are shaded in gray. The position of the LXCXE RB1 binding site is highlighted. A cartoon of the domain structure of E7 is shown on top, with the N-terminal sequences that show similarity the conserved regions 1 and 2 (CR1, CR2) of adenovirus E1A proteins indicated **(A)**. MmuPV1 E7 (M E7) can interact with RB1 by immunoprecipitation/immunoblot analysis. Duplicate cultures of HCT116 cells were transfected with an N-terminally HA/FLAG epitope-tagged MmuPV1 E7 expression plasmid and co-precipitated RB1 protein detected by immunoblotting **(B)**. MmuPV1 E7 (M E7) can interact with murine Rb1 by immunoprecipitation/immunoblot analysis. NIH 3T3 murine fibroblasts were transfected with an N-terminally HA/FLAG epitope-tagged MmuPV1 E7 expression plasmid and co-precipitated RB1 protein detected by immunoblotting **(C)**. MmuPV1 E7 does not destabilize RB. SAOS-2 human osteosarcoma cells that do not express detectable endogenous RB were transfected with an RB expression vector and increasing amounts of an expression vector for N-terminally HA/FLAG epitope-tagged MmuPV1 followed by western blotting to assess RB1 steady levels by western blotting. GFP was co-transfected and assessed by western blotting to control for transfection efficiency. HPV16 E7 (16 E7) was used as a positive control. A representative blot from one of four experiments is shown **(D)**.

The canine papillomavirus 2 (CPV2) E7 protein, which, similar to MmuPV1 E7, lacks an LXCXE domain, has been reported not only to bind RB1 but also to destabilize it (16). Given the importance of RB1 destabilization by high-risk HPVs in cellular transformation (22), we asked whether MmuPV1 E7 can destabilize RB1. RB1 was co-transfected in combination with increasing amounts of MmuPV1 E7 into SAOS-2 human osteosarcoma cells, which express an inactive, C-terminally truncated, barely detectable RB1 mutant (23). In contrast to HPV16 E7 (22, 24), expression of MmuPV1 E7 did not cause a significant decrease in RB1 steady-state levels. Hence MmuPV1 does not cause detectable RB1 destabilization (**Figure 2D**).

### MmuPV1 E7 does not as efficiently activate E2F-dependent gene expression as HPV16 E7

One of the best-studied biological activities of RB1 is its regulation of the activity of the transcription factor E2F1 (25–27). RB1 undergoes cell cycle-dependent phosphorylation and dephosphorylation. Hypophosphorylated RB1 binds to E2F family members and the resulting RB1/E2F transcriptional repressor complexes restrain transition from the G1 to the S phase of the cell cycle. When RB1 is hyperphosphorylated by cyclin-dependent kinases, it permits E2Fs to function as transcriptional activators and drive S-phase progression (28, 29). Like adenovirus E1A and polyomavirus large tumor antigens, LXCXE motif-containing HPV E7 proteins bind RB1 and abrogate the formation of RB1/E2F repressor complexes thereby causing aberrant S-phase entry. It is thought that this activity of the viral proteins is key to retaining infected host cells in a replication-competent state that is conducive for viral genome synthesis (30). To determine whether MmuPV1 binding to RB1 affects E2F transcription factor activity, we determined expression levels of four well-established, E2F-regulated genes, cyclin E2 (CCNE2) (31), minichromosome maintenance complex components 2 (MCM2) and 7 (MCM7) (32) as well as Proliferating Cell Nuclear Antigen (PCNA) (33) in MmuPV1 E7 expressing, telomerase-immortalized human keratinocytes (iHFKs) or early passage mouse keratinocytes. HPV16 E7 expressing cells were used as controls. While expression of CCNE2, MCM2, MCM7, and PCNA was significantly increased in HPV16 E7-expressing iHFKs, there was no comparable increase in the expression of these genes in MmuPV1 E7-expressing iHFKs (**Figure 3A**) or primary mouse keratinocytes (**Figure 3B**). Based on these results, we conclude that MmuPV1 E7 does not as efficiently activate the expression of E2F-responsive genes as HPV16 E7.

**Figure 3.**
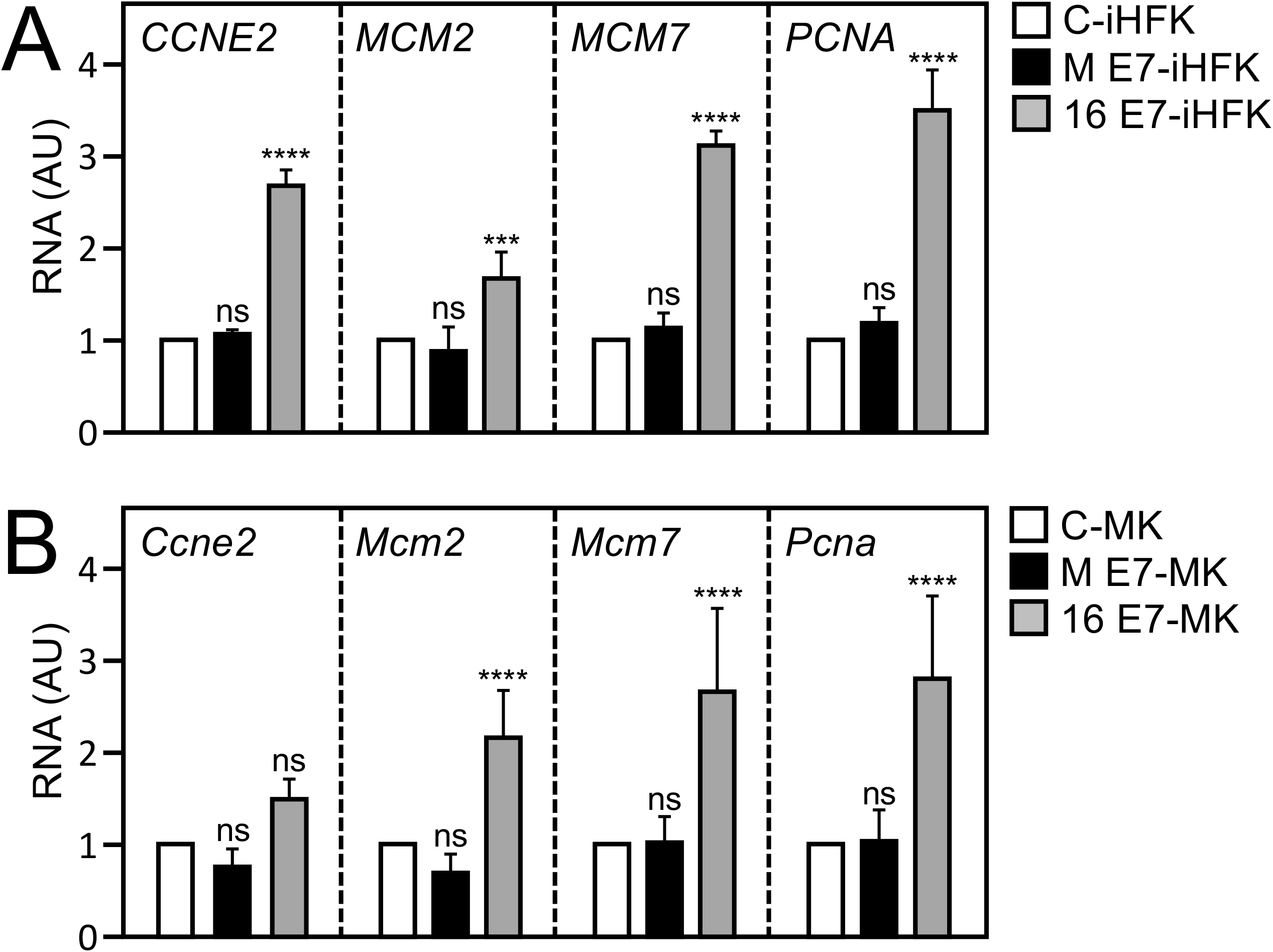
MmuPV1 E7 does not efficiently activate E2F-regulated genes. Expression of E2F target genes, cyclin E2 (CCNE2), Minichromosome Maintenance Complex Components 2 and 7 (MCM2, MCM7) and Proliferating Cell Nuclear Antigen (PCNA) in immortalized human foreskin keratinocytes (iHFKs) **(A),** or in primary mouse keratinocytes (MK) **(B)** transduced with control vector (C), MmuPV1 E7 (M E7) or HPV16 E7 (16 E7) as determined by quantitative reverse transcription-coupled polymerase chain reaction analysis (****, *p*<0.001; ***, *p*<0.005; n.s. not significant).

### The MmuPV1 E7 protein interacts with RB1 sequences that are distinct from the LXCXE binding cleft

Because MmuPV1 E7 does not contain an LXCXE sequence, we wanted to test whether it binds RB1 similarly or distinctly to LXCXE-containing E7 proteins. The LXCXE binding site in RB1 has been determined by X-ray co-crystallography studies (34). Based on this information, an LXCXE binding defective RB1 mutant with amino acid substitutions at three critical contact residues (I753A; N757A; M761A) (RB1^L^) was constructed (35). We compared the abilities of HPV16 E7, which binds RB1 through its LXCXE motif, and MmuPV1 E7 to bind to wild-type RB1 versus the RB1^L^ mutant by expressing the corresponding proteins in SAOS-2 cells. As expected (35), HPV16 E7 interacted with wild-type RB1 but not the RB1^L^ mutant. In contrast, MmuPV1 interacted with both wild-type RB1 and mutant RB1^L^ with similar efficiencies. These experiments reveal that MmuPV1 interacts with RB1 sequences that are distinct from those necessary for interaction with LXCXE motif-containing E7 proteins (**Figure 4**).

**Figure 4.**
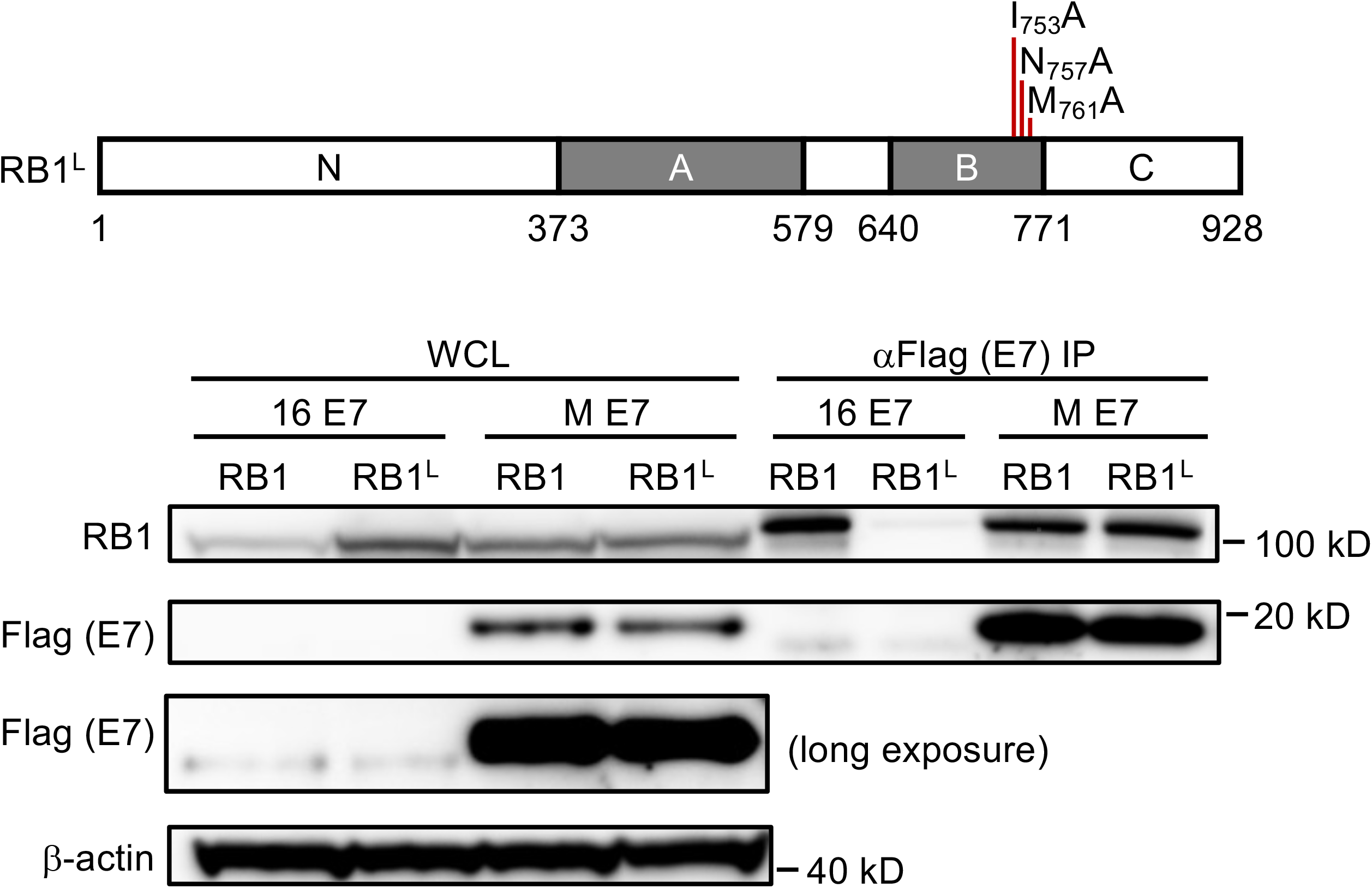
The LXCXE binding cleft in the RB1 protein is not necessary for MmuPV1 E7 binding. SAOS-2 human osteosarcoma cells were transfected with expression vectors for wildtype RB1 or the RB1^L^ mutant that contains three amino acid mutations in the LXCXE binding cleft in combination with MmuPV1 or HPV16 E7 expression vectors. MmuPV1 E7 binds wild type and RB1^L^ with similar efficiency whereas HPV16 E7 binds and causes degradation of RB1 but not RB1^L^. The results shown are representative of two independent experiments. A cartoon of the RB1^L^ mutant is shown at the top of the figure.

### MmuPV1 causes warts in mice expressing the LXCXE binding defective Rb1^L^ mutant

We previously used *Rb1^L^* knock-in mice expressing the above described mutant Rb1 to investigate the role of HPV16 E7’s binding to Rb1 in neoplastic disease (36, 37). Given that MmuPV1 E7 can interact with the RB1^L^, we asked if MmuPV1 can cause disease in *Rb1^L^* mice. Ears of both wild-type *FVB* and *Rb1^L^ FVB* mice were infected with 10^8^ VGE of MmuPV1 (virus stock generated from MmuPV1-induced warts from a nude mouse - see Materials and Methods) as described previously (20) using the same methodology used in our previous experiments for quasivirus infections. By the end of 4 months, MmuPV1 caused a similar incidence of papillomas in both the wild type and *Rb1^L^ FVB* mice (**Figure 5A**, *p-value* = 1, two-sided Fisher’s exact test). There was no apparent size difference between warts from the MmuPV1-infected wild-type mice and those from *Rb1^L^* mice, based on H&E analysis (data not shown).

**Figure 5.**
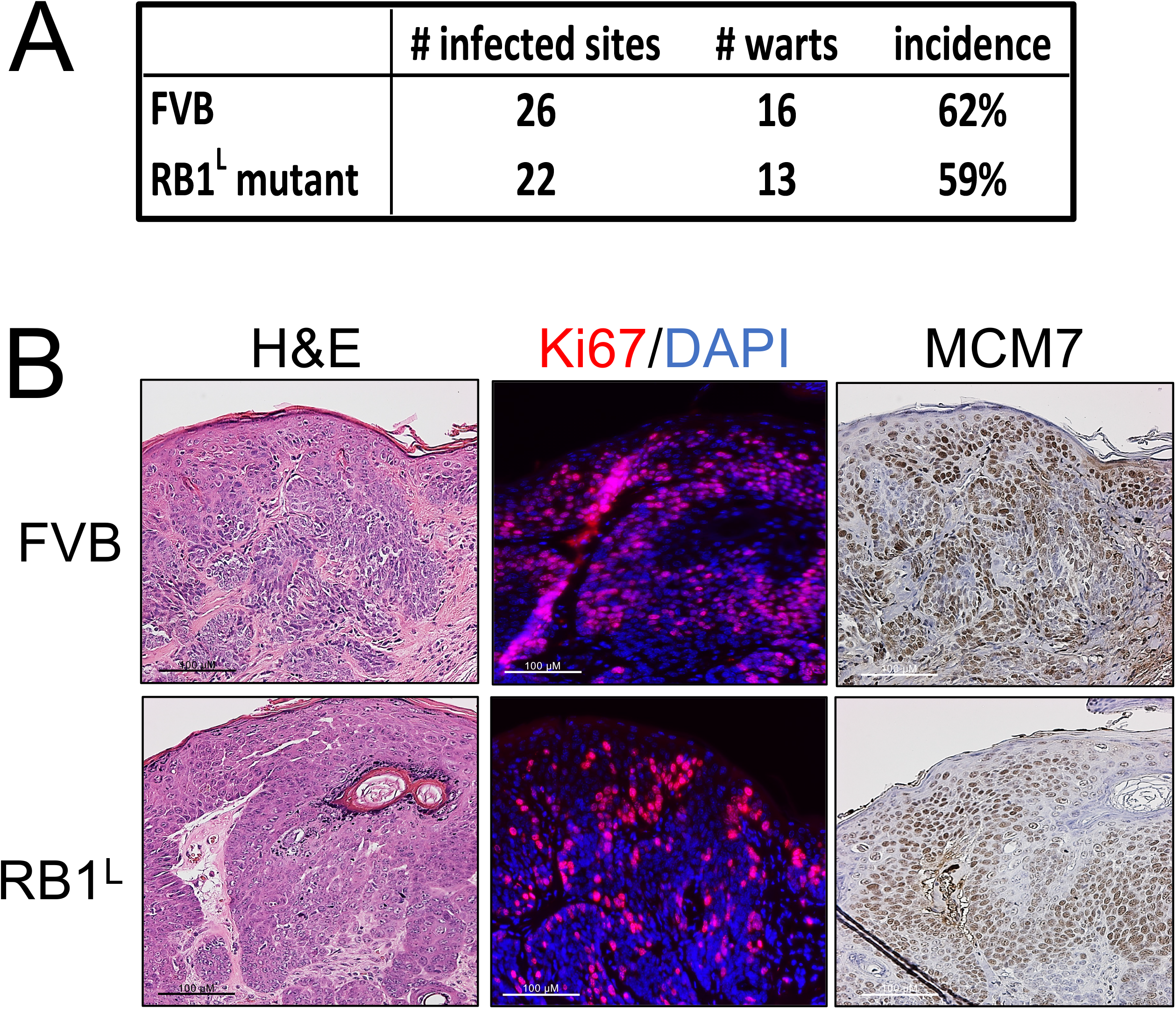
Disruption of the LXCXE binding cleft in RB1 does not influence MmuPV?s ability to cause papillomas in vivo. Sites on the ears of both wild-type and RB1^L^ mutant FVB/N mice were scarified and infected with 10^8^ VGE of MmuPV1. Mice were treated with 300 mJ UVB the next day and then monitored for papilloma formation over 4 months. MmuPV1 induced warts in wild type and RB1^L^ mutant FVB/N mice with a similar incidence (Fisher’s exact test, p Value=1, two-sided) **(A)**. Warts arising in wild-type and RB1^L^ mutant FVB/N mice share similar microscopic features. Warts from both mouse genotypes were harvested, serially sectioned, and stained with hematoxylin and eosin (H&E), processed to detect Ki67 (Ki67, red; DAPI, blue) by immunofluorescence, and MCM7 by immunohistochemistry **(B)**.

We next performed immunohistochemistry on tissues obtained at the experimental endpoint to compare the expression of biomarkers for viral infection between the papillomas arising in wild type and *Rb1^L^ FVB* mice (**Figure 5B**). Ki67-specific immunohistochemistry showed similar levels of cell proliferation in the papillomas arising on both the wild type and *Rb1^L^* FVB mice. MCM7 was also upregulated to similar levels, indicating that MmuPV1 is capable of increasing E2F-driven gene expression despite the disruption in the LXCXE binding cleft in the *Rb1^L^ FVB* mice. These results demonstrate that the disruption of the ability of RB1 to bind proteins via their LXCXE-motifs is not required for MmuPV1 to induce papillomas or to induce expression of E2F-responsive genes.

### C-terminal RB1 sequences are necessary for interaction with MmuPV1 E7

Given that MmuPV1 E7 does not interact with the LXCXE binding cleft of RB1 we mapped the RB1 region responsible for binding MmuPV1 E7. We co-expressed MmuPV1 E7 with plasmids expressing full-length RB1 (amino acid residues 1-928) or truncation mutants of RB1 lacking the amino-terminus (amino acid residues 379-928) or the C-terminus (amino acid residues 1-792) in SAOS2 human osteosarcoma cells. HPV16 E7 was used as a control. As expected, HPV16 E7 efficiently bound wild-type RB1, as well as the two truncation mutants, both of which encode the A and B domains of RB1 that contain the LXCXE binding cleft. In contrast, MmuPV1 E7 did not efficiently interact with the 1-792 mutant RB1 that lacks the C-terminal domain of RB1 (**Figure 6A, B**). Hence MmuPV1 E7 primarily interacts with the C-terminal domain of RB1.

**Figure 6.**
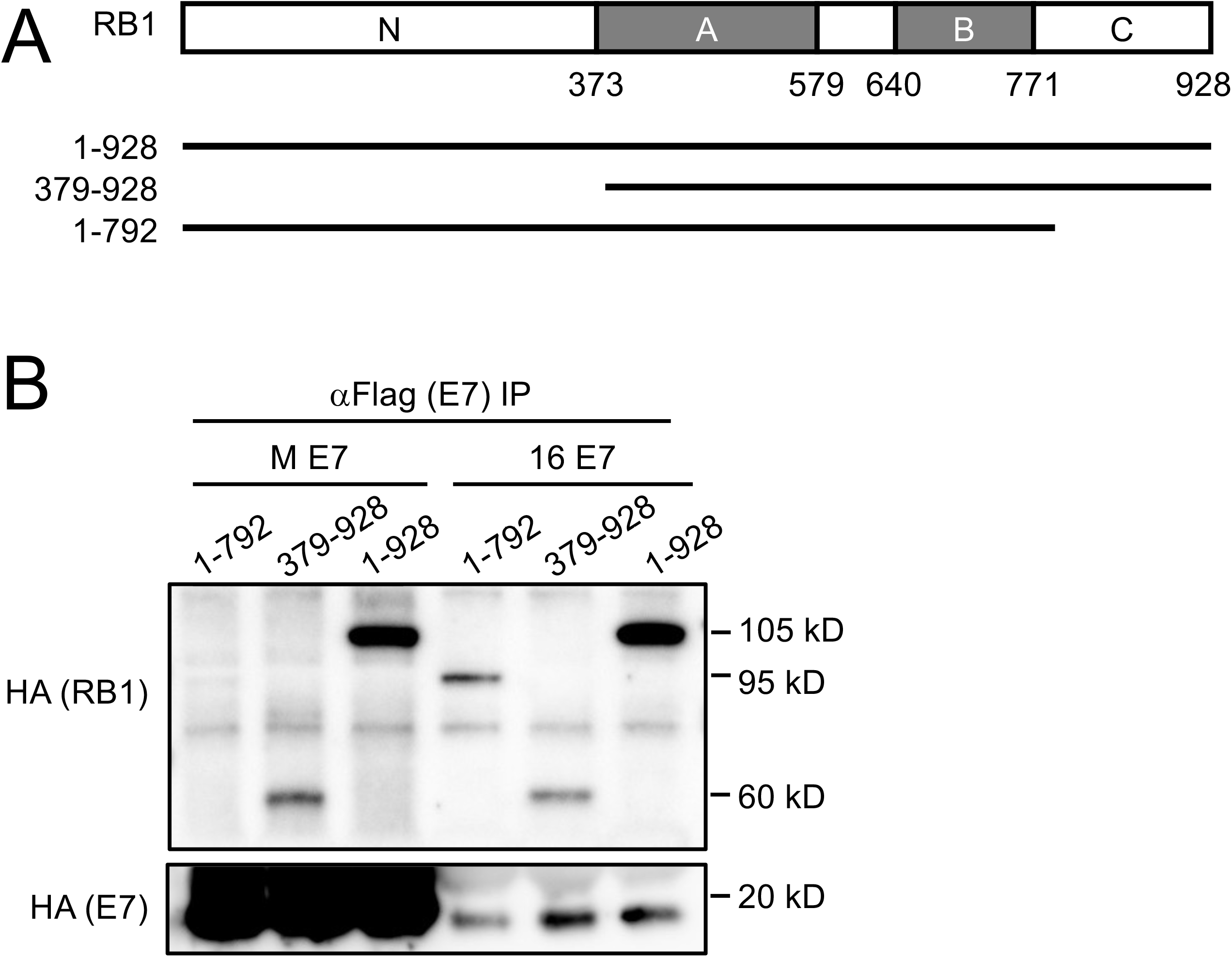
The C-terminal domain of RB1 is necessary for MmuPV1 E7 binding. Schematic representation of the RB1 protein and the expression plasmids used **(A)**. HA-epitope tagged versions of full-length RB1, and the two truncation mutants 1-792 and 379-928 were expressed in SAOS-2 human osteosarcoma cells in combination with FLAG/HA epitope-tagged MmuPV1 (M E7) and HPV16 E7 (16 E7) expression vectors. After immunoprecipitation with FLAG antibodies, HA-tagged E7 and co-precipitated RB1 proteins were detected by HA immunoblot. The result shown is representative of two independent experiments. **(B)**.

### C-terminal MmuPV1 E7 sequences are necessary for RB1 binding

It has previously been reported that the E7 proteins of CPV2 and HPV4, which also lack LXCXE motifs in their N-termini, associate with RB1 through their C-termini (16). Hence, in addition to testing some mutations in the N-terminus of MmuPV1 E7, we also generated MmuPV1 E7 mutants in the C-terminal domain. We focused on regions that are conserved between CPV2, HPV4, HPV197, and MmuPV1 (**Figure 7A)**. Of all the mutants that were tested, a four amino acid deletion of residues 84 to 87 (MmuPV1 E7^Δ84-87^) and an alanine substitution at aspartate residue 90 (MmuPV1 E7^D90A^) were found to be defective for RB1 binding (**Figure 7B**). Based on these results we generated two additional substitution mutants, MmuPV1 E7^D90T^ and MmuPV1 E7^D90N^. MmuPV1 E7^D90T^ was generated to mimic the HPV16 E7 threonine residue at this position (**Figure 7A**). MmuPV1 E7^D90T^ displayed decreased RB1 binding similar to MmuPV1 E7^D90A^. The MmuPV1 E7^D90N^ mutant, which was generated to neutralize the negative charge while maintaining the general architecture of the side chain, retained some binding to RB1, even though it was expressed at lower levels than the other two mutants **(Figure 7C)**. Lastly, the MmuPV1 E7^D90A^ mutant was also defective for binding to murine Rb1 **(Figure 7D)**. Hence, similar to CPV2 and gamma-HPVs (16), MmuPV1 E7 binds to RB1 through its C-terminal domain, and based on the results described above, we chose the MmuPV1 E7^D90A^ mutant for our follow-up studies.

**Figure 7.**
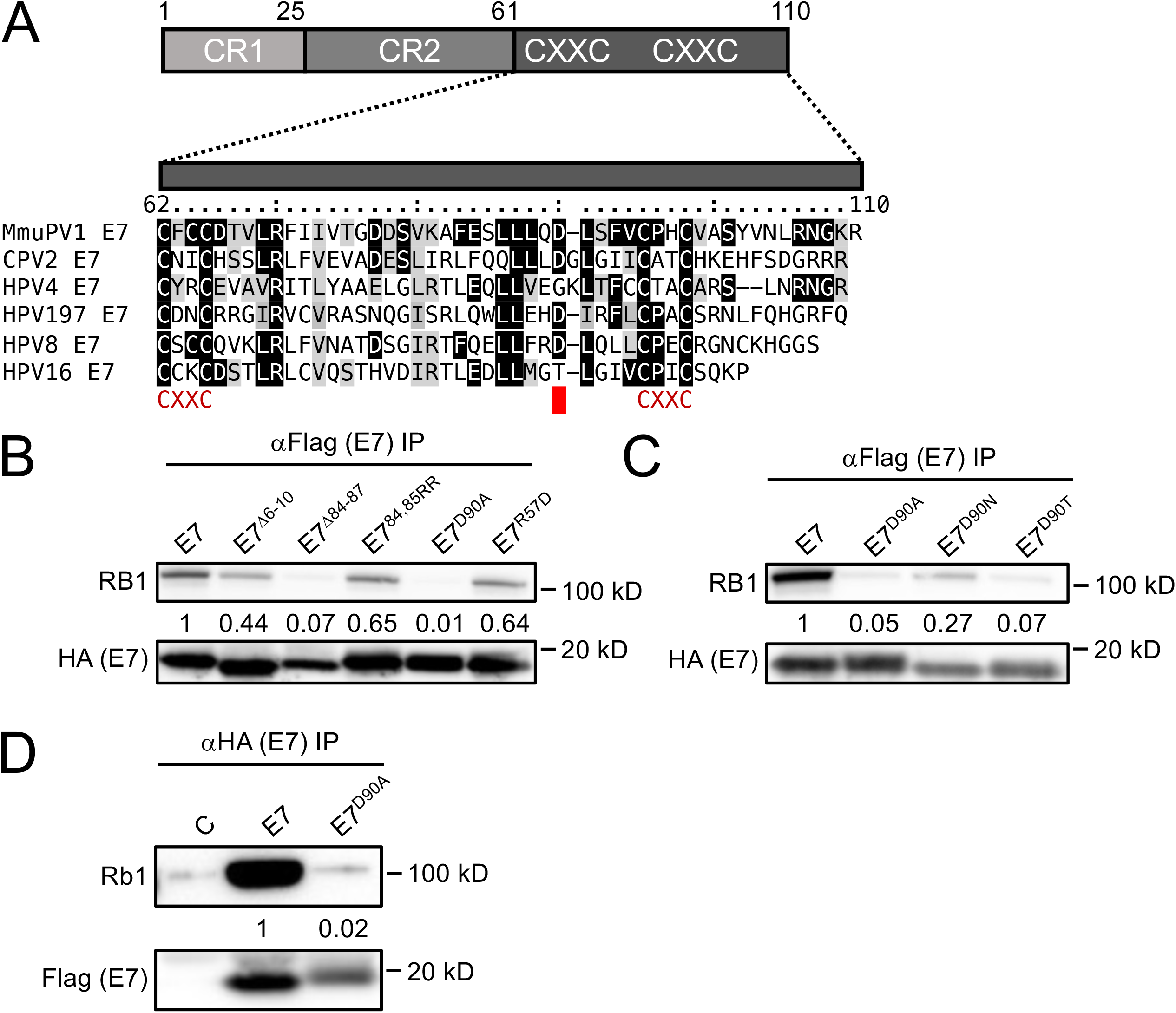
The RB1 binding site maps the MmuPV1 E7 C-terminus. Sequence alignment of the C-terminal domains of MmuPV1, canine papillomavirus 2 (CPV2), γ1-HPV4, γ24-HPV197, β1-HPV8, and α9-HPV16 E7. Identical residues are marked by black boxes, chemically similar residues are shaded in gray. The position of the CXXC motifs that form a zinc-binding site is shown. The position of the aspartate residue at position 90 (D90) that is important for RB1 binding is indicated by a red box **(A)**. Various FLAG/HA-tagged MmuPV1 E7 mutants were expressed in HCT116 cells and co-precipitated RB1 was detected by immunoblotting. The result shown is representative of six independent experiments **(B)**. Immunoprecipitation western blot analyses to assess RB1 binding by various MmuPV1 E7^D90^ mutants. The result shown is representative of two independent experiments **(C)**. Immunoprecipitation/western blot analysis documenting that MmuPV1 E7^D90A^ is defective for binding to murine Rb1. Wild type MmuPV1 E7 and the D90A mutant were transiently expressed in NIH3T3 cells and binding assessed by immunoprecipitation/western blotting. The blot shown is representative of 2 independent experiments **(D)**. Quantifications of E7-coprecipitated RB1 or Rb1 are normalized to the amount of E7 that is precioitated and are shown underneath.

### Reduced incidence and smaller sized warts in MmuPV1 E7^D90A^ infected mice

To assess whether E7’s ability to bind Rb1 contributes to MmuPV1 pathogenesis, we introduced the D90A mutation into the complete MmuPV1 DNA genome which we used to make quasivirus particles in 293FT cells. We then characterized the ability of this mutant MmuPV1 versus wild-type MmuPV1 to cause papillomatosis in mice. Quasivirus stocks were confirmed to be infectious (**Figure 8A**), by exposing mouse keratinocytes to the quasiviruses and 48 hours later harvesting RNA to detect the presence of viral E1^E4 spliced transcripts by RT-PCR. We then performed *in vivo* infections with these stocks of infectious quasivirus. 6-8 weeks old Nude-*FoxN1^nu/nu^* mice were scarified on their ears and tails and infected with wildtype MmuPV1 or E7^D90A^ mutant MmuPV1 quasivirus at doses of either 10^7^ (stock 2) or 10^8^ VGE (stock 1). Papilloma incidence was monitored biweekly for 4 months. At the endpoint, wild-type MmuPV1 at the 10^8^ VGE dose caused papillomas at 100% of the sites infected. In contrast, the same dose of MmuPV1 E7^D90A^ quasivirus caused papillomas at a significantly lower frequency, 12% (**Figure 8B**, p<0.0001, two-sided Fisher’s exact test). At the lower 10^7^ VGE dose, wild-type MmuPV1 caused papillomas at a 65% frequency, whereas MmuPV1 E7^D90A^ caused papillomas in only 8% of infected sites (**Figure 8B**, p<0.001, two-sided Fisher’s exact test). At both doses, the papillomas arising on mice infecting with MmuPV1 E7^D90A^ appeared at later time points (**Figure 8C**, MmuPV1 (10^8^ VGE) vs. MmuPV1 E7^D90A^ stock 1, p<0.0001; MmuPV1(10^7^ VGE) versus MmuPV1 E7^D90A^ stock 2, p<0.0001 two-sided LogRank test). At the 4 month endpoint, we harvested the papillomas from all infected mice, fixed and serially sectioned the lesion, and performed H&E staining. We performed scans of the H&E-stained sections from 6 representative papillomas induced by MmuPV1 (3 from 10^8^ VGE and 3 from 10^7^ VGE), and 3 papillomas induced by MmuPV1 E7^D90A^ quasiviruses (we scanned all three papillomas that arose on mice infected with the MmuPV1 E7^D90A^ quasivirus, regardless of virus dose) (**Figure 8D**). The size of each papilloma was assessed using ImageScope under the same magnification (**Figure 8E**). Papillomas caused by the MmuPV1 E7^D90A^ quasivirus were significantly smaller when compared to those caused by MmuPV1 (MmuPV1 versus MmuPV1 E7^D90A^, p= 0.003, two-sided T-test), indicating that the loss of E7’s ability to interact with Rb1 correlates with reduced size of MmuPV1-induced papillomas. We harvested and sequenced the MmuPV1 genomes present in warts arising from mice infected with the wild type and the E7^D90A^ quasiviruses and confirmed that warts indeed contained the expected virus and that no cross-contamination had occurred. Together, the assessments of the incidence of papillomas, the time of onset of papillomas, and the size of papillomas at the endpoint all indicate that the interaction between MmuPV1 E7 and Rb1 quantitatively correlates with the MmuPV1’s ability to cause papillomatosis.

**Figure 8.**
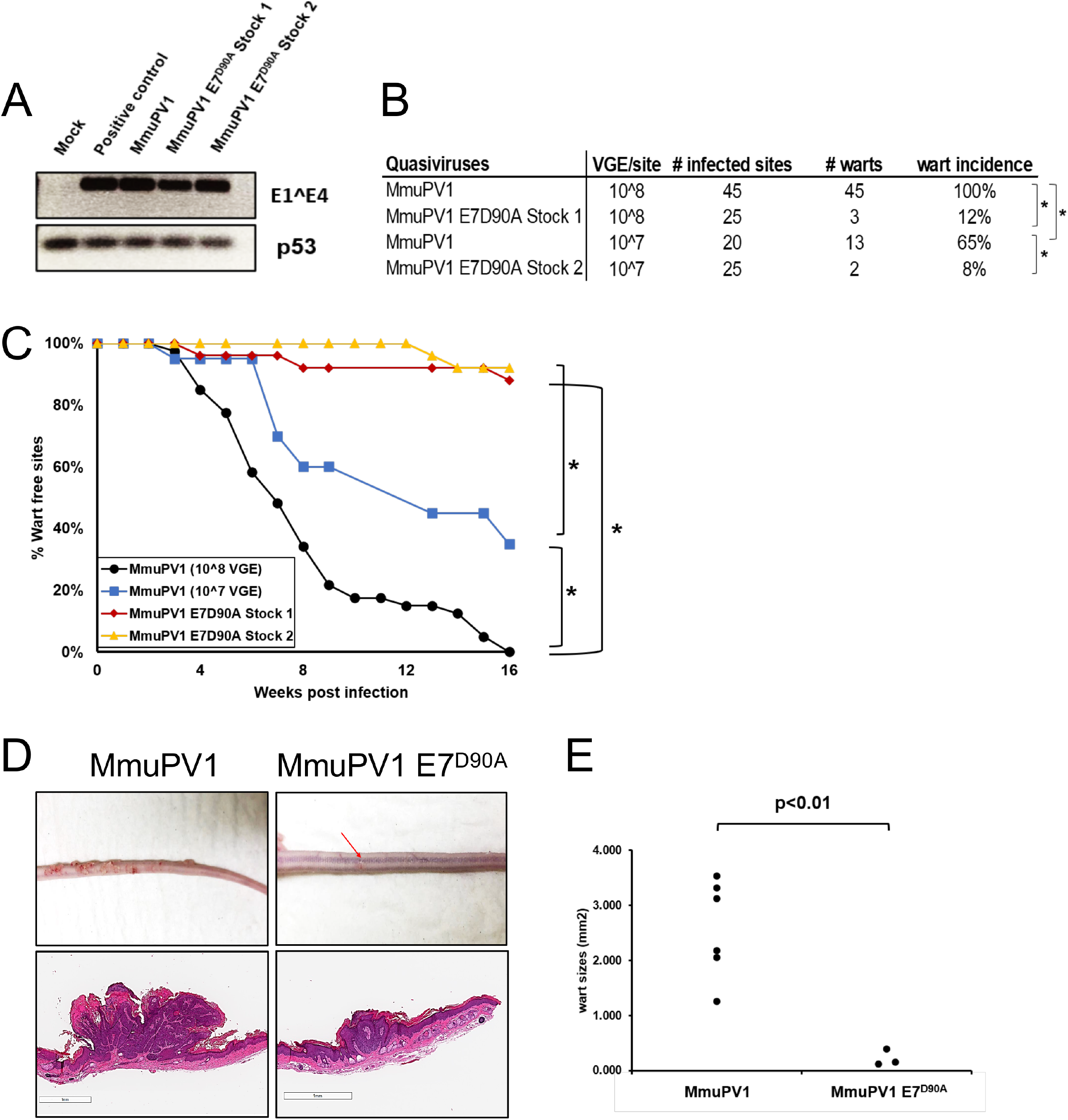
MmuPV1 E7^D90A^ virus gives rise to reduced incidence and smaller-sized warts than wild type MmuPV1. To assess infectivity of quasivirus stocks, mouse JB6 keratinocytes were exposed to quasivirus and 48 hours later RNA was extracted and subjected to reverse transcription-coupled polymerase chain reaction (RT-PCR) to detect MmuPV1 E1^E4 (top panel) or p53 (bottom panel) transcripts. Shown are results for mock-infected cells, and cells infected with equal amounts of MmuPV1 derived from warts (positive control), wild type MmuPV1 quasivirus (wild type), and two stocks of E7^D90A^ mutant quasivirus (E7^D90A^) **(A)**. Wart incidence arising at sites on nude mice infected with 10^8^ VGE of wild type MmuPV1 quasivirus, 107 VGE of wild type MmuPV1 quasivirus as well as 10^8^ or 10^7^ VGE, respectively, of two independent preparations of MmuPV1 E7^D90A^ quasivirus (E7^D90A^ Stock 1, E7^D90A^ Stock 2). The incidence of warts at sites infected with MmuPV1 E7^D90A^ quasivirus was significantly less compared to dose equivalent of wild type MmuPV1 quasivirus (Fisher’s exact test, two-sided: MmuPV1 (10^8^ VGE) vs. E7^D90A^ Stock 1, p<0.0001; MmuPV1 (10^7^ VGE) vs. E7^D90A^ Stock 2, p<0.001). A dosage effect in wart formation was also observed with the wildtype MmuPV1 (MmuPV1 (10^8^ VGE) vs. MmuPV1 (10^7^ VGE), p=0.0001) **(B)**. Kaplan-Meier plot showing that the percent of wart free sites over time is significantly different for MmuPV1 versus MmuPV1 E7^D90A^ quasivirus infections (LogRank test (two-sided): MmuPV1 (10^8^ VGE) vs. MmuPV1 E7^D90A^ Stock 1, p<0.0001; MmuPV1 (10^7^ VGE) versus MmuPV1 E7^D90A^ Stock 2, p<0.0001; MmuPV1 (10^8^ VGE) versus MmuPV1 (10^7^ VGE), p<0.001) **(C)**. Representative images and of tails of mice infected with the different quasiviruses at the 4 months endpoint (top images), along with equal magnification of scanned images of sections of tails harboring representative warts stained with H&E (bottom images) **(D)**. Size of warts at the 4 months endpoint. Warts arising from the MmuPV1 E7^D90A^ quasivirus were significantly smaller (T-test, two-sided: MmuPV1 versus MmuPV1 E7^D90A^, p=0.003). Six MmuPV1 warts (three from each dose) and all three MmuPV1 E7^D90A^ quasivirus induced warts (two from Stock 1 and one from Stock 2) were used for this quantification.

### MmuPV1 E7^D90A^ - induced papillomas display similar histological features as papillomas induced by wild-type MmuPV1

To determine whether the papillomas caused by MmuPV1 E7^D90A^ displayed similar or different microscopic features compared to those caused by wild-type MmuPV1, we performed immunohistochemistry to assess the expression of biomarkers for papillomavirus-associated lesions. Evidence for productive viral infections within the papillomas was scored by performing immunofluorescence staining to detect the viral capsid protein L1 (**Figure 9**, panel B). L1 expression was similar in papillomas induced by the wild type and mutant quasiviruses. Papillomas induced by wild type and E7^D90A^ quasiviruses also showed similar patterns of keratinocyte differentiation with cytokeratin 14 upregulated in the suprabasal layers of the papillomas (**Figure 9**, panel B), indicating similar delays in terminal differentiation. MCM7, an E2F-responsive gene that is upregulated in papillomavirus-related lesions caused by high-risk HPV (38) and MmuPV1 (39) infections, was similarly upregulated in papillomas induced by wild type and E7^D90A^ quasiviruses, indicating increased levels of E2F-mediated transcription in both cases (Figure 9, panel C). The incorporation of BrdU into genomic DNA is often upregulated in papillomavirus-related lesions as a consequence of increased DNA synthesis. Both MmuPV1 and MmuPV1 E7^D90A^ - induced papillomas showed increased levels of BrdU, with no obvious differences in abundance or localization of BrdU-positive cells within the papillomas, indicating similarly enhanced levels of DNA synthesis (**Figure 9**, panel D). Based on the biomarkers tested, there were no significant differences in the histopathological features due to the loss of interaction between E7 and Rb1.

**Figure 9.**
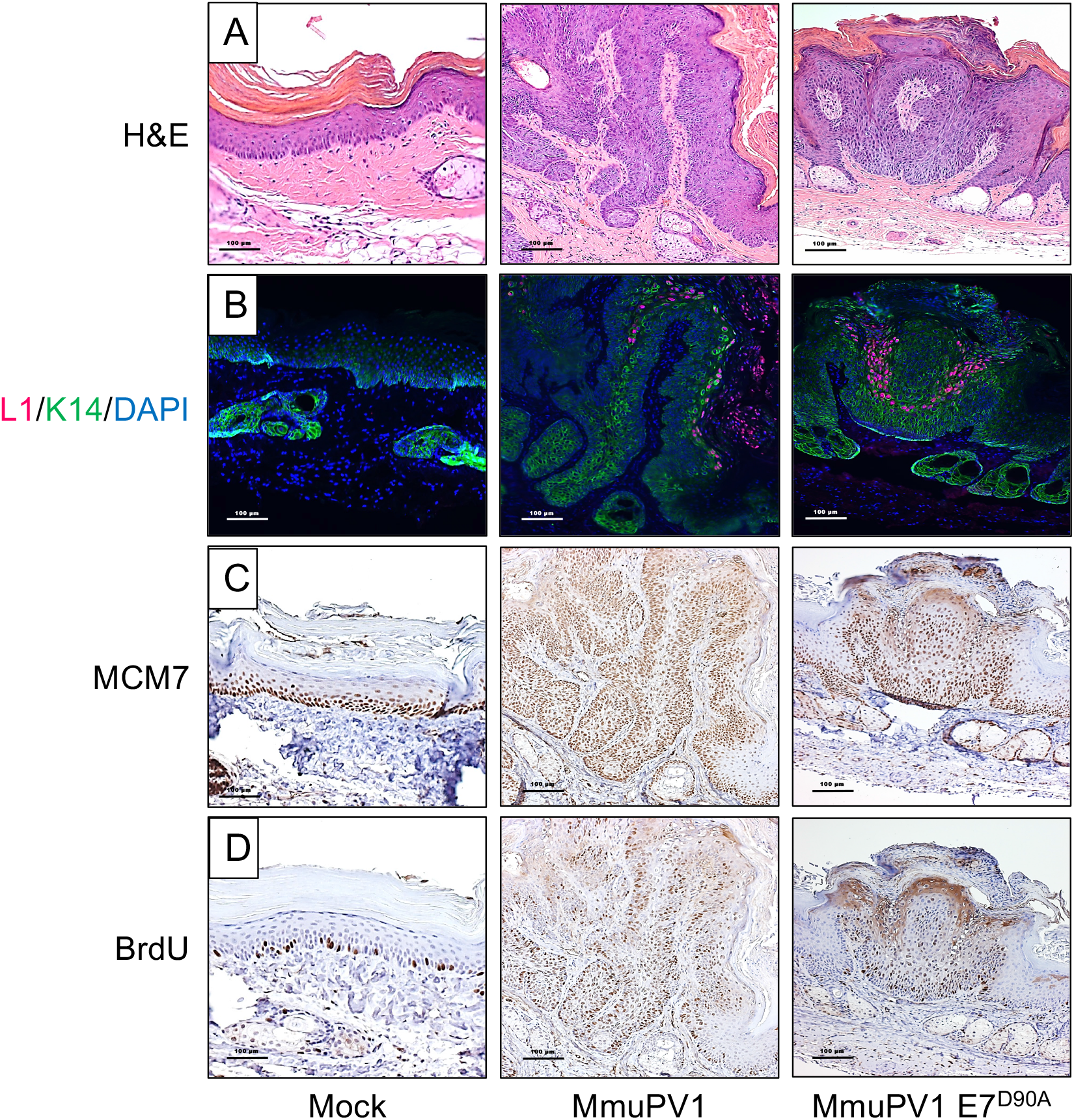
MmuPV1 E7^D90A^ quasivirus-induced warts display similar histological features as warts induced by wildtype MmuPV1. Shown are serial sections of the mock-infected tail (left column), and tail warts induced by wildtype MmuPV1 (middle column) or MmuPV1 E7^D90A^ quasivirus (right column) stained with H&E **(A)**, stained for MmuPV1 L1 (red), K14 (green) by immunofluorescence and counterstained with DAPI (blue) **(B)**, or immunohistochemically stained for BrdU **(C)**, or MCM7 **(D)**.

## DISCUSSION

The species specificity of papillomaviruses has greatly limited studies of how specific biochemical activities of individual viral proteins contribute to viral pathogenesis in a natural infection model. The discovery of MmuPV1 and its ability to infect and cause papillomas in laboratory mice has removed this barrier. MmuPV1 is a member of the Pi genus, which encompasses rodent PVs (1) and is most related to the cutaneous beta- and gamma-genus HPVs (40).

The MmuPV1 E6 and E7 proteins share sequence similarities to cutaneous beta- and gamma-HPVs, respectively (41). We have previously reported that MmuPV1 E6 is necessary for papilloma formation in experimentally infected mice (15). MmuPV1 E6 shares key cellular targets and biological activities with the beta-HPVs 5 and 8 E6 proteins that affect key tumor suppressor gene functions, including the ability to bind the NOTCH transcriptional coactivator MAML1 and the SMAD2 and SMAD3 mediators of transforming growth factor-beta (TGF-beta) signaling (15). In particular, we have shown that MmuPV1 E6’s ability to bind NOTCH correlates with MmuPV1’s ability to cause disease (15).

Here we show that similar to what we previously reported for E6 (15), MmuPV1 E7 is also necessary for papilloma formation. Like CPV2 E7 and some gamma-HPV E7 proteins, the MmuPV1 E7 protein lacks an N-terminal LXCXE motif, which is present in multiple viral and cellular proteins where it serves as the binding site for members of the retinoblastoma tumor suppressor family (40). We provide evidence that MmuPV1 E7 can bind to both the human and murine retinoblastoma tumor suppressor proteins. We determined that the RB1 binding site is located in the MmuPV1 E7 C-terminal domain, similar to CPV2 and gamma-HPV4 E7 proteins (16). We identified a C-terminal mutant, MmuPV E7^D90A^, that was markedly reduced for RB1 binding. The HPV16 E7 C-terminus may also contain a low-affinity RB1 binding site (42) and mutation of the T86 residue in HPV16 E7 (which corresponds to MmuPV1 E7 D90) to an aspartate (as present in MmyPV1) did not significantly affect RB1 binding in a yeast two-hybrid format (43), whereas the MmuPV1 E7 E7^D90T^ mutant exhibited decreased RB1 binding. Infection with the MmuPV1 E7^D90A^ mutant revealed that RB1 binding correlates with MmuPV1’s ability to promote efficient papilloma formation in the cutaneous infection model. However, it does not appear to be essential for pathogenesis as small warts did arise, albeit at significantly reduced efficiency and with a later time of onset. There is precedence for these findings as studies that were done in cottontail rabbit PV (SfPV1, a.k.a. CRPV1) found E7 to be essential for promoting disease (44, 45) but that E7’s ability to interact with RB1, albeit through an LXCXE sequence (46), is not essential for inducing papillomas in experimentally infected rabbits (47). In addition, studies with transgenic mice have also provided evidence that HPV16 E7 can cause hyperproliferation in mice that express the *Rb1^L^* mutant, which HPV16 E7 is unable to bind (36, 37). Therefore, it is likely that other biological activities of PV E7 are also playing important roles in promoting disease across various PVs.

PV E7 proteins with an LXCXE motif bind to a shallow cleft within the RB1 B “pocket” domain (34). In contrast, MmuPV1 E7 interacts with the RB1 C-terminal domain. Consistent with these results, experimental MmuPV1 infection of mice engineered to express the *Rb1^L^* allele, in which the LXCXE binding cleft in the B domain is mutated, caused papilloma formation at a similar efficiency as mice infected with wild type MmuPV1 genomes. The RB1 C-terminal domain is highly conserved amongst RB1 proteins from different species and is necessary for RB1 to induce permanent G1 growth arrest and senescence (48–50). It has been shown to mediate interactions with several different cellular proteins, including the ABL1 non-receptor tyrosine kinase (51), the F-box protein SKP2 (52) and a non-canonical E2F1 complex that contains the lysine methyltransferase EZH2 (53, 54). It has been reported that ABL1 selectively binds to hypophosphorylated RB1 and that RB1 binding inhibits ABL1 enzymatic activity (51). While the RB1 C-terminus is necessary, the ABL1 binding sequences have not been mapped in detail and the biological relevance of the RB1/ABL1 interaction has remained enigmatic. SKP2 is an F-box protein that is part of the cullin1 based ubiquitin ligase complex that has been shown to control the degradation of the CDK2 inhibitor, p27^KIP1^ (CDKN1B). SKP2 is rapidly degraded during G1, when RB1 is hyperphosphorylated, by CDH1 containing anaphase-promoting complex or cyclosome (APC/C^CDH1^) (55) that binds to RB1’s A-B pocket. However, when RB1 gets hyperphosphorylated, APC/C^CDH1^ dissociates from RB1 which leads to SKP2 degradation of p27^KIP1^ and increased CDK2 activity which promotes S-phase entry. These and other non-canonical RB1 activities (56) including the ability of RB1 to interact with E2F1/EZH2 complexes that are not involved in cell cycle regulation are likely targeted by MmuPV1 E7, but further studies are necessary to identify the specific target(s) and to delineate the molecular consequences of C-terminal RB1 binding.

Given that phosphorylation-specific RB1 binding and release of E2F transcription factor complexes involve RB1 sequences within the A-B domain (57, 58), and that MmuPV1 E7 interacts with RB1 C-terminus, it was not surprising that MmuPV1 E7 expression did not cause efficient activation of E2F-responsive genes. Nevertheless, MmuPV1 can induce expression of MCM7, a strongly E2F-responsive gene, in vivo in the context of papillomas it induces. We also observed MCM7 induction in MmuPV-1-induced papillomas arising in mice expressing the *Rb1^L^* allele and in papillomas arising in animals infected with MmuPV1 genomes expressing the RB1 binding defective E7^D90A^ mutant. This raises the interesting question: which MmuPV1 protein(s) is responsible for the increased MCM7 expression. Experiments performed with HPVs in cell and transgenic animal-based models have all suggested that this activity is provided by E7 and is based on its ability to inactivate RB family members and it is possible that E7^D90A^ mutant retains low-level RB1 binding that may be sufficient to cause some expression of E2F-responsive genes in vivo. However, our study is not to first to document E7 independent induction of hyperproliferation. Experimental infection of rabbits with an SfPV1 mutant virus that expressed an RB binding-deficient E7 mutant still caused the emergence of papillomas (47). Our work is entirely consistent with this observation; moreover, our studies provide no evidence that MmuPV1 E7 can efficiently activate the expression of E2F-responsive genes. It is possible that MmuPV1 encodes another protein that can activate E2F-dependent promoters through direct or indirect mechanisms. MmuPV1 E6 has been hypothesized to be able to bind Rb1 (59) because it contains an LXCXE motif (L_67_ACKE_71_) in between its two (CXXC)_2_ zinc-binding domains; however, our prior AP/MS experiments failed to provide any evidence that MmuPV1 E6 binds to any of the RB family members (13). Another possibility is that the expression of E2F-responsive genes in the papillomas reflects the ability of MmuPV1 E6 to impede keratinocyte differentiation through its inhibition of NOTCH and TGF-beta signaling which may help MmuPV1 infected cells maintain a proliferative state (15). Regardless, given that MmuPV1 E7 expression is necessary for papilloma formation, it will be important to determine the mechanism by which MmuPV1 E7 contributes to papilloma formation. Infections with an RB1 binding defective E7 mutant gives rise to smaller papillomas with lower efficiency and delayed kinetics compared to papillomas caused by wild-type MmuPV1 infection. It will be important to determine whether papillomas that express the RB1 binding defective E7 mutant progress to cancer at a similar frequency, or at all, compared to papillomas caused by wild-type MmuPV1.

In summary, our results show that loss of RB1 binding by MmuPV1 E7 correlates with a quantitative defect in papilloma induction. Hence MmuPV1 E7 binding to RB1’s C-terminal domain remains an important mechanism by which MmuPV1 promotes disease. The integrity of the RB1 C-terminus is important for many activities of RB1, but whether or how any of these contribute to Rb1’s tumor suppressor activity is largely unknown. Given, the vast majority of studies on RB1 have focused on its ability to control E2F transcription factor activity, which is shared with other RB family members that are not frequently mutated in tumors, it is unlikely that regulation of E2F transcription factor activity is the sole tumor-suppressive function of RB1. It will be important to rigorously determine which specific function of RB1’s C-terminus MmuPV1E7 disrupts. Such studies promise to provide exciting new insights into the molecular basis of RB1 ‘s tumor suppressor activity.

## MATERIALS AND METHODS

### Cells

U2OS human osteosarcoma cells were obtained from ATCC and grown in Dulbecco’s Modified Eagle Medium (DMEM; Invitrogen) supplemented with 10% fetal bovine serum (FBS). SAOS-2 human osteosarcoma cells were obtained from ATCC and grown in McCoy’s 5A medium (Invitrogen) supplemented with 15% FBS. HCT116 human colon carcinoma cells were obtained from ATCC and grown in McCoy’s 5A medium (Invitrogen) supplemented with 10% FBS. NIH 3T3 murine fibroblasts were obtained from the ATCC and grown in DMEM (Invitrogen) containing 5% FBS. JB6 mouse keratinocytes (gift from Dr. Nancy H. Colburn, NCI) were maintained in DMEM containing 5%FBS. 293FT cells (ATCC) were maintained in DMEM with 10%FBS and 300 μg/mL neomycin (G418). Telomerase-immortalized human foreskin keratinocytes Cl398 (iHFK) (60) were a kind gift from Aloysius Klingelhutz (U. of Iowa.). iHFK lines expressing various E7 proteins were established by transducing iHFKs with the corresponding pLenti-NmE7 expressing lentiviruses, followed by selecting with 3 μg/ml Blasticidin (RPI Research Products International, Mount Prospect, IL) for 7 days starting at two days post-infection. Mouse Keratinocytes were isolated from the skin of neonate pups. After incubation in PBS containing 10% antibiotics for 2 minutes, skin pieces were incubated in 0.25% trypsin overnight at 4°C. The epidermis was then separated from the dermis using sterile forceps, minced with a single edge razor blade, and then stirred for 1 hour at 37°C in F-media (61) to generate a single-cell suspension. The cells were strained using 0.7 μm membrane (102095-534; VWR), and cultured in F-media (62) containing 10 μM Y-27632 Rho-kinase inhibitor (63) in the presence of mitomycin C (M4287; Sigma) treated 3T3 J2 fibroblasts. Early passage cells were infected with the recombinant lentiviral or retroviral vectors expressing HPV16 E7 or MmuPV1 E7, respectively, in F-medium in the absence of Y-27632 and 3T3 J2 feeders and re-infected after 24 hours. At 72 hours after the first infection, cells were selected with the appropriate antibiotics. After selection, cells were maintained in F-media containing 10 μM Y-27632 and mitomycin C-treated 3T3 J2 fibroblasts.

### Plasmids and antibodies

The MmuPV1 E7 ORF was PCR amplified from the MmuPV1 genome and cloned into N- or C-terminal FLAG/HA-CMV (64) and untagged pCMV-BamNeo (65) plasmids. Mutant MmuPV1 E7 constructs were generated by site-directed mutagenesis of N-FLAG/HA-mE7-CMV. pLenti-N-mE7 was generated by Gateway cloning (Invitrogen) of PCR-amplified mE7 into pLenti 6.3/V5 DEST (Invitrogen). The RB1-truncation plasmids pSG5-HA-Rb 1-928, 1-792, and 379-928 were kind gifts from Bill Sellers (Broad Institute). The pFADRb and pFADRbL plasmids were kindly provided by Fred Dick (Western University, Ontario). Other plasmids used were pLXSN HPV16 E7 (66), pCMV-Rb (67) (obtained from Phil Hinds, Tufts), CMV-C-16E7 (17), pBABE puro (68), and pEGFP-C1 (Clontech). The following primary antibodies were used for immunoprecipitations and western blotting: beta-Actin (MAB1501; Millipore), FLAG (F3165; Sigma), GFP (9996; Santa Cruz), HA (ab9110; Abcam), RB1 (Ab-5, OP66; Millipore), and Rb1 (SC-74570, Santa Cruz). Secondary anti-mouse or anti-rabbit antibodies conjugated to horseradish peroxidase were from GE Healthcare.

### Immunological Methods

Affinity purification/mass spectrometry analyses of MmuPV E7 were performed as previously described (17). HCT116 cells were transfected using Polyethylenimine (PEI) (69). At 48 hr post-transfection cells were harvested in EBC buffer (50mM Tris-Cl pH 8.0, 150 mM NaCl, 0.5% NP-40 and 0.5mM EDTA) supplemented with protease inhibitors (Pierce). Anti-Hemagglutinin (HA; Sigma) or anti-Flag epitope (Sigma) antibodies coupled to agarose beads were used for immunoprecipitations followed by SDS-PAGE and western blot analysis on PVDF membranes. After incubation with appropriate primary and secondary antibodies, blots were visualized by enhanced chemiluminescence and images captured using a Syngene ChemiXX6 imager with Genesys software version 1.5.5.0. Signals were quantified with Genetools software version 4.03.05.0.

### RB1 Degradation assays

RB1 degradation assays were performed as previously described (22). SAOS-2 cells were transfected with CMV-RB and varying amounts of pCMV N-FLAG/HA mE7. pCMV-C16 E7 was used as positive control and pEGFP-C1 was co-transfected to assess transfection efficiency. At 48 hours post-transfection cells were lysed in EBC and samples containing 100 μg protein were subjected to western blot analysis as described above.

### Quantitative Reverse Transcription PCR

RNA was isolated from pLenti-N-FLAG/HA mE7, iHFK pLenti-C-FLAG/HA-16E7, and pLenti-N-GFP infected iHFKs and pLenti-N-FLAG/HA mE7, pLXSP-16E7 or control vector infected primary mouse keratinocytes with the Quick-RNA Miniprep kit (Zymo Research). cDNA was synthesized with the Quantinova Reverse Transcription Kit (Qiagen). Quantitative PCR was performed in triplicate on a Step One Plus (Applied Biosystems) thermocycler using Fast SYBR Green Master Mix (Applied Biosystems). PCR primers are listed in supplemental Table S2. Target expression levels were normalized to GAPDH expression.

### Animals

Immunodeficient Athymic *Nude-FoxN1^nu/nu^* mice were purchased from Envigo. *RB1^L^* mutant mice were maintained on the FVB background and genotyped as published previously (37, 70). Mice were housed in the Association for Assessment of Laboratory Animal Care-approved McArdle Laboratory Animal Care Unit. All procedures were carried out in accordance with an animal protocol approved by the University of Wisconsin Institutional Animal Care and Use Committee (IACUC; protocol number M005871).

### Infection of nude mice with MmuPV1 quasiviruses

MmuPV1 quasiviruses (wild type, E7^STOP^, E7^D90A^; note that the term “quasivirus” is used in the papillomavirus field to identify a virus that is generated in cells by co-transfection of viral genomes with a plasmid that expresses the viral capsid proteins) were generated as described before (15). Briefly, 293FT cells were transfected with a MmuPV1 capsid protein expression plasmid (71, 72) and the re-circularized genome, either from plasmids containing the wild type or mutant MmuPV1 genomes. After incubation at 37 °C for 48 hours, cells were harvested and virus extracted. The amount of packaged viral DNA in the stocks of quasiviruses was quantified by Southern blotting allowing us to define the ‘viral genome equivalents” (VGE), as a measure of virus concentration in each stock. The quasiviruses were used to infect Nude-*FoxN1^nu/nu^* mice as previously described (15). Briefly, animals were placed under anesthesia and infected by first scarifying the epidermis using a 27-gauge syringe needle and then pipetting onto the wounded site the indicated amount of quasivirus using a siliconized pipette tip. Each mouse was infected at 5 sites maximum (one site per ear, three sites on tail). Papillomatosis was monitored weekly/ bi-weekly as indicated.

### Infection of FVB-background mice with MmuPV1

The infection method has been described previously with some modifications (20). Briefly, under anesthesia, mouse ears of both FVB and *RB1^L^* mutant mice were scarified first using 27-gauge syringe needles and infected with 10^8^ VGE/site of a pre-prepared stock of MmuPV1 virus generated from a MmuPV1-infected wart. 24 hours later, mice were exposed to 300mJ UVB (Daavlin, Bryan, OH). Papillomatosis was monitored over 4 months.

### RT-PCR to detect MmuPV1 E1^E4 spliced transcripts

Mouse keratinocytes JB6 cells were infected with MmuPV1 wild type and mutant quasiviruses at 10^8^ VGE, and changed to fresh media 3 hours later. After incubation at 37°C for 48 hours, total RNA was extracted from infected JB6 cells using the RNeasy kit (Qiagen) and reverse-transcribed into cDNA using the QuantiTect Reverse Transcription Kit (Qiagen). E1^E4 transcripts were detected by PCR, using p53 as a positive control. Primer sequences were described previously (72).

### BrdU incorporation

To evaluate levels of DNA synthesis, we performed bromodeoxyuridine (BrdU) incorporation by injecting BrdU (Sigma, dissolved in PBS to 12 mg/ml stock concentration, keep at −20°C). Mice were intraperitoneally injected with 250 μl stock BrdU one hour before harvest. Tissues were harvest and processed for immunohistochemistry using a BrdU-specific antibody (203806, Calbiochem) as previously described (15).

### Histological analysis

Tissues were harvested and fixed in 4% paraformaldehyde (in PBS) for 24 hours, then switched to 70% ethanol for 24 hours, processed, embedded in paraffin, and sectioned at 5 μm intervals. Every 10th section was stained with hematoxylin and eosin (H&E).

### MmuPV1 L1-cytokeratin dual immunofluorescence and Immunohistochemistry

L1 signals were detected using a tyramide-based signal amplification (TSA) method (73). A detailed protocol is available at: https://www.protocols.io/view/untitled-protocol-i8cchsw.

For immunohistochemistry, tissue sections were deparaffinized in xylenes and rehydrated in 100%, 95%, 70%, and 50% ethanol, then in water. Antigen unmasking was performed by heating with 10 mM citrate buffer (pH=6) for 20 minutes. Blocking was performed with 2.5% horse serum in PBST for 1 hour at room temperature (RT). Slides were incubated in primary antibody (BrdU; MCM7, Thermo Scientific, Fremont, CA) at 4°C, overnight in a humidified chamber. M.O.M.® ImmPRESS® HRP (Peroxidase) polymer kit (Vector, MP-2400) was applied the next day for 1 hour at RT for secondary antibody incubation. Slides were then incubated with 3,3’-diaminobenzidine (Vector Laboratories), and counterstained with hematoxylin. All images were taken with a Zeiss AxioImager M2 microscope using AxioVision software version 4.8.2.

### Full scan for wart size measurement, and statistical analysis

Full scans of representative warts were performed by the UW Translational Research Initiatives in Pathology (TRIP) facility. Measurements were performed on the full-scanned images using ImageScope software (v12.4.0). All statistical analyses were performed using MSTAT statistical software version 6.4.2 (http://www.mcardle.wisc.edu/mstat).

## ACKNOWLEDGMENTS

We thank Dr. Al Klingelhutz (University of Iowa) for providing telomerase immortalized human foreskin keratinocytes, members of the Lambert and Munger labs for stimulating discussion and valuable suggestions throughout the course of this work, Ella Ward-Shaw for expert histotechnology assistance, and Simon Blaine-Sauer for isolating C57/B6 mouse keratinocytes from neonates. Supported by PHS grant R01 CA228543 (K.M. and P.F.L.) and a Ruth L. Kirschstein Postdoctoral Individual National Research Service Award F32 CA254019 (J.R.).

## Table Legends

**Table S1:** Cellular proteins identified by Affinity purification/mass spectrometry (AP/MS) analyses of carboxyl- or amino terminally epitope tagged MmuPV1 E7 (CE7 and NE7 respectively). The number of unique and total peptides identified for each protein are shown. See reference [17] for experimental details.

**Table S2: Table S2:** Sequences of PCR primers used in this study.

